# Structures of the LPOR–Chlide Complexes Imply the Basis of Membrane Remodeling and a Photocatalytic Mechanism

**DOI:** 10.64898/2025.12.12.693892

**Authors:** Michał Gabruk, Ambroise Desfosses, Leandro Farias Estrozi, Sebastian Pintscher, Michał Rawski, Grzegorz Ważny, Agnieszka Garbacz, Mateusz Zbyradowski, Jerzy Kruk, Leszek Fiedor

## Abstract

Flowering plants rely on the photocatalytic enzyme light-dependent protochlorophyllide oxidoreductase (LPOR) to synthesize chlorophyllide (Chlide), a chlorophyll intermediate, and at the same time to remodel lipid membranes into the cubic phase required for chloroplast development. Yet neither the mechanism of this light-driven catalysis nor the structural basis of its membrane remodeling activity is well understood, largely due to the lack of high-resolution structural information.

To address these questions, we analyzed Chlide:LPOR:NADPH oligomeric assemblies by cryo-electron microscopy. Eleven maps were obtained, enabling the reconstruction of nine distinct LPOR oligomer models. Most assemblies adopt helical or stacked-ring forms, whereas one displays a segmented string-of-dimers architecture, representing a previously unobserved structural organization.

Strings of LPOR dimers induce distinct periodic deformations of the lipid bilayer that stabilize multiple complex architectures. These architectures are maintained by three different inter-string interfaces occurring in various combinations. The plasticity of these interactions suggest that the filaments are capable of twisting and sliding relative to each other. Our highest-resolution map (2.55 Å) provided a detailed view of the pigment-binding site, revealing a water channel that connects the central magnesium ion of the pigment to the bulk solvent. Unexpectedly, the propionate group of the pigment protrudes out of the porphyrin plane and arches back toward NADPH, ideally positioned for hydride transfer.

These structural insights allowed us to propose a novel reaction mechanism for LPOR photocatalysis. Together, our data highlight the critical role of Chlide:LPOR:NADPH complexes in delaying prolamellar body disassembly and point to their possible regulatory function in mature leaves.

## Introduction

Flowering plants rely on the photocatalytic enzyme, named light-dependent protochlorophyllide oxidoreductase (LPOR), to synthesize a chlorophyll (Chl) intermediate in a light-dependent manner and at the same time to remodel lipid membranes into a cubic phase required for chloroplast development. Yet neither the mechanism of this light-driven catalysis nor the basis of membrane remodeling activity is well understood, largely due to a lack of high-resolution structural information.

It has been shown that the longer seedlings grow in darkness, the more LPOR accumulates in ternary complexes with NADPH and the penultimate intermediate of the Chl biosynthetic pathway^1^, protochlorophyllide (Pchlide), in immature chloroplasts (etioplasts). These complexes remodel the inner membranes of etioplasts into a paracrystalline lattice, the prolamellar body (PLB), whose main lipid component is the non-bilayer-forming monogalactosyldiacylglycerol (MGDG)^2–5^.

Thus far, a few crystal structures of cyanobacterial LPOR have been determined ^6,7^, however, these isoforms do not remodel membranes into cubic phases in vitro^8^, and the structures lacked bound pigment. However, the cryo-EM structure of the oligomeric Pchlide:LPOR:NADPH from *A. thaliana* at 3.2 Å resolution^9^ revealed the binding pocket and showed that the enzyme forms helical assemblies composed of strings of dimers that trap a constricted bilayer inside the filamentous complexes. Similar assemblies were observed within PLBs by cryo-tomography^10^. The structure revealed two parts of the pigment-binding pocket—helix α10 and the Pchlide loop—embedded in the outer leaflet, with an MGDG molecule bound next to the pigment^9^. The lipid binding redshifts the pigment emission to ∼655 nm at 77 K ^11^, characteristic of PLB. However, in vitro–generated complexes lack the cubic ultrastructure, which can be induced by adding NADP⁺ first to a reaction mixture containing LPOR, Pchlide, and a lipid mixture mimicking the native PLB composition^8^, followed by the addition of NADPH to displace NADP⁺ and form the photoactive complex within the cubic phase^8^.

After exposure to light, LPOR uses NADPH to reduce one double C-C bond of the tetrapyrrole ring of the substrate, converting Pchlide to chlorophyllide (Chlide). The photoreaction is triggered by absorption of a photon by the pigment, but there is no consensus on the subsequent events. Some recent QM/MM studies^12,13^ modeled alternative substrate orientation rather than the cryo-EM–based pose to remain consistent with earlier mechanistic models; other studies employed the cryoEM-based orientation and pointed to Y306 as a proton donor^14^, in contrast to earlier proposals suggesting Y276 based on homology to other family members^15,16^.

Regardless of the mechanism, it has been shown that immediately after the illumination the Chlide emission initially redshifts, which some researchers interpreted as NADPH rebinding to the Chlide:LPOR complex after photoconversion^17,18^. A similar reaction cycle was proposed for the cyanobacterial LPOR^16^. After prolonged illumination, the emission blueshifts, in a process known as the Shibata shift^19^, interpreted as disassembly of LPOR oligomers^20–24^, which is associated with subsequent esterification of Chlide, that completes Chl synthesis. Interestingly, ATP was shown to accelerate the Shibata shift^25^, what ultimately leads to PLB disassembly and, consequently, thylakoid formation^26^.

Despite decades of research, the mechanisms of the photoreaction and of ultrastructural reorganization during PLB formation and disassembly remain unclear. Therefore, we investigated the events occurring after illumination of native PLB, supported by parallel experiments in an in vitro system. As a result, we determined the structures of several LPOR assemblies with Chlide, revealing novel details of the binding site, proposing new reaction mechanism of the photoreaction and membrane remodeling, and pointing to the role of Chlide:LPOR complexes in mature leaves.

## Results

### Chlide:LPOR:NADPH complex formation delays PLB disassembly

To investigate the molecular events triggered by illumination of the PLB we used ∼2 cm-long segments of etiolated wheat seedlings and fluorescence spectroscopy. To capture ultrafast events, we applied a cryostat set to 80 K and monitored changes in the fluorescence emission spectrum of these pigments as the temperature increased (Fig. 1A, bottom; Extended Data Fig.1AB). This temperature-resolved approach enables detection of successive reaction steps as they become thermally accessible. At 80 K, an emission peak at 655 nm, characteristic of PLB, was observed, which originates from active ternary Pchlide:LPOR:NADPH. Upon heating, the signal at 655 nm diminished almost entirely by 160 K, with nearly no detectable emission from Chlide at that point (Fig. 1A bottom, Extended Data Fig.1AB). Chlide fluorescence became detectable only above 160 K, with a maximum around 687 nm, resembling the spectrum of in vitro reconstituted Chlide:LPOR:NADP⁺ complexes (Fig.1C, Extended Data Fig.1C). At temperatures exceeding 260 K, the emission peak blue-shifted to ∼681 nm, which resembles emissions of Chlide:LPOR complexes lacking the dinucleotide cofactor in vitro (Fig.1C, Extended Data Fig.1C).

**Figure 1.**
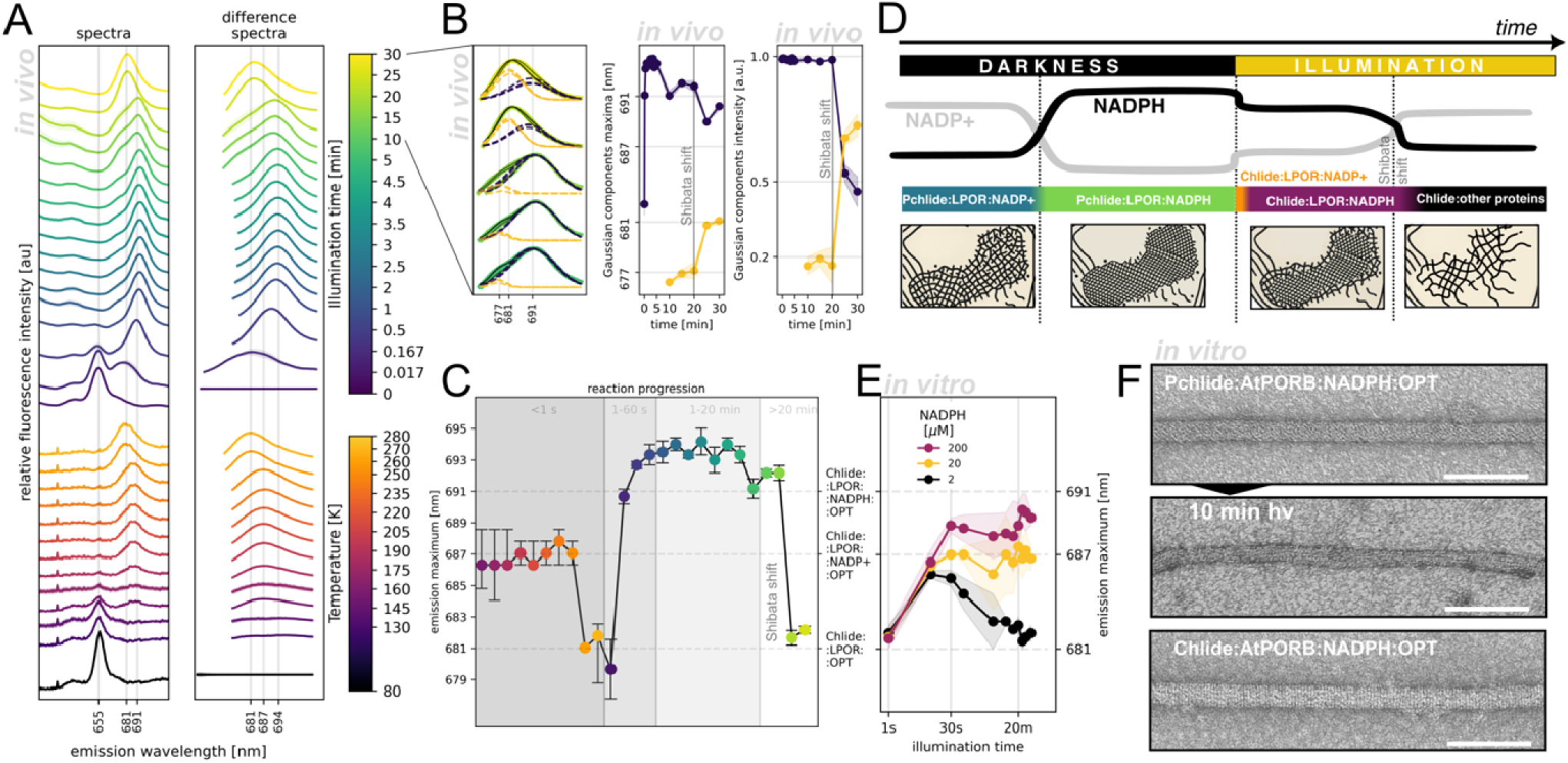
Chlide:LPOR:NADPH oligomeric complexes form in PLB after illumination. **A.** Fluorescence spectra of wheat seedlings illuminated over a temperature range (bottom) or for varying time intervals and measured at 77 K (top). Shaded areas represent standard deviation between biological replicates. **B.** Gaussian deconvolution of selected spectra, showing emission maxima and intensities; shaded areas represent standard deviation between biological replicates. **C.** Relationship between Chlide emission maximum and reaction progress, calculated from the data in A. **D.** Model of PLB disassembly inferred from shifts in Chlide emission, showing relative NADPH and NADP⁺ concentrations, the composition of LPOR complexes, and a schematic cross-section of the PLB ultrastructure. **E.** Time-course of Chlide emission maximum versus illumination time for in vitro reaction mixtures containing recombinant LPOR with varying NADPH concentrations (15 µM AtPORB, 5 µM Pchlide, 40 µM OPT lipids, and NADPH). Shaded area represent standard deviation between three replicates. **F.** Negative-stain transmission electron micrographs of reaction mixtures (15 µM AtPORB, 5 µM pigment, 40 µM OPT lipids, 200 µM NADPH) incubated in darkness or illuminated; scale bar = 100 nm.

To further probe the subsequent molecular events, we again used fragments of etiolated wheat seedlings and illuminated them at room temperature, but the reaction was stopped by rapid freezing in liquid nitrogen at various time intervals. To better visualize the signal originating from the reaction product, we calculated difference spectra by subtracting the average spectrum of the unilluminated sample and then fitted one or two Gaussian curves (Fig. 1A top, Extended Data Fig. 1D). We observed that the product of the photoreaction appeared already after 1 second of illumination, with a fluorescence maximum at ∼679 nm, which is consistent with Chlide:LPOR complexes devoid of the dinucleotide. Apparently, this method is too slow to detect the initial NADP⁺ complex, suggesting that the post-reaction state somehow rapidly triggers the dinucleotide release. After additional 10 seconds of illumination, the emission maximum red-shifted to 690-695 nm and remained at that position for several minutes. This range of emission maximum resembles that of Chlide:LPOR:NADPH complexes reconstituted in vitro in the presence of MGDG-containing membranes (OPT lipids: 50mol% MGDG, 35mol% DGDG, 15mol% PG; Extended Data Fig.1C). This suggests that, after Chlide formation and the release of NADP⁺, NADPH rebinds to the active site within 20 seconds after the illumination.

A prolonged incubation of illuminated etiolated seedlings led to an additional blueshift of Chlide emission, known as Shibata shift. Analysis of the Gaussian components of Chlide emission during this phase revealed that the peak consists of two distinct components: one at around 690 nm, which indicates Chlide:LPOR:NADPH complexes, while the other at 677-681 nm (Fig.1C). During the Shibata shift, their relative intensities reversed (Fig. 1C), resulting in the shift of the emission from 690 to 681 nm. Although the resulting Chlide emission maximum resembles that of Chlide:LPOR complex lacking the dinucleotide, it is plausible that similar emission originates from the complexes of the pigment with some other proteins present within etioplasts.

These observations suggest that, after illumination, PLB disassembly is delayed due to the formation of Chlide:LPOR:NADPH complexes (Fig. 1D). To test this hypothesis, we conducted in vitro kinetic measurements using varying NADPH concentrations and observed a clear correlation: higher NADPH concentrations resulted in a more pronounced red shift of the fluorescence emission maximum (Fig. 1F), which is consistent with formation of Chlide:LPOR:NADPH complexes. Finally, to assess whether these complexes can form stable oligomers, we used transmission electron microscopy (TEM) (Fig. 1E). We found that filamentous LPOR oligomers indeed persisted even after 10 minutes of illumination at high NADPH concentrations. Moreover, the periodic oligomeric complexes could be reconstituted in vitro by adding purified Chlide directly to LPOR in the presence of OPT lipids and NADPH. These observations strongly support the notion that LPOR in vivo forms stable oligomers not only with Pchlide but also with Chlide if NADPH and lipids containing MGDG are present. To elucidate how LPOR accommodates both Pchlide and Chlide, we performed helical reconstruction of Chlide:AtPORB:NADPH:OPT complexes imaged by cryo-EM.

### LPOR assembles into a variety of oligomeric states

Within a single dataset collected from one grid (18 466 videos, 40 frames each), we observed a diversity of filamentous assemblies, varying in both architecture and diameter (Fig. 2ABC, Extended Data Fig. 2, Extended data video 1, Extended Data Table 1-9). Despite this heterogeneity, we successfully classified seven distinct subsets, each enabling three-dimensional reconstructions at resolutions ranging from 2.9 to 3.7 Å (Fig. 2D). All assemblies were composed of LPOR dimers organized into strings that surrounded a tubular membrane bilayer. Three of these assemblies adopted a helical architecture, with diameters of 230 Å, 250 Å, and 290 Å, therefore we designated them as **H**elical **F**ilaments HF-23, HF-25, and HF-29, respectively (Fig. 2D). Assemblies HF-23 and HF-25 were composed of four strings of dimers (“4-start” helices), while HF-29 consisted of three strings (“3-start” helix). Another assembly resembled a helical architecture but was segmented into three-subunit-long fragments arranged as a single string of dimers (“1-start” helix). Since it was 250 Å in diameter, we named it SF-25 (**S**egmented **F**ilament). Three other assemblies exhibited a stacked-ring architecture with diameters of 210 Å, 230 Å, and 250 Å, and were thus named **R**ing **F**ilaments RF-21, RF-23, and RF-25, respectively. These ring-like assemblies had resolution of 3.2 Å and were composed of 10, 11, and 12 LPOR dimers per ring. To improve the resolution of the dimers composing each stacked-ring architectures, we performed sub-particle averaging of all dimeric units, yielding maps at 2.6–2.7 Å resolution, which we designated RD-21, RD-23, and RD-25, respectively. We then merged these datasets to produce a final map at 2.55 Å resolution, termed RD-A (Fig. 2D).

**Figure 2.**
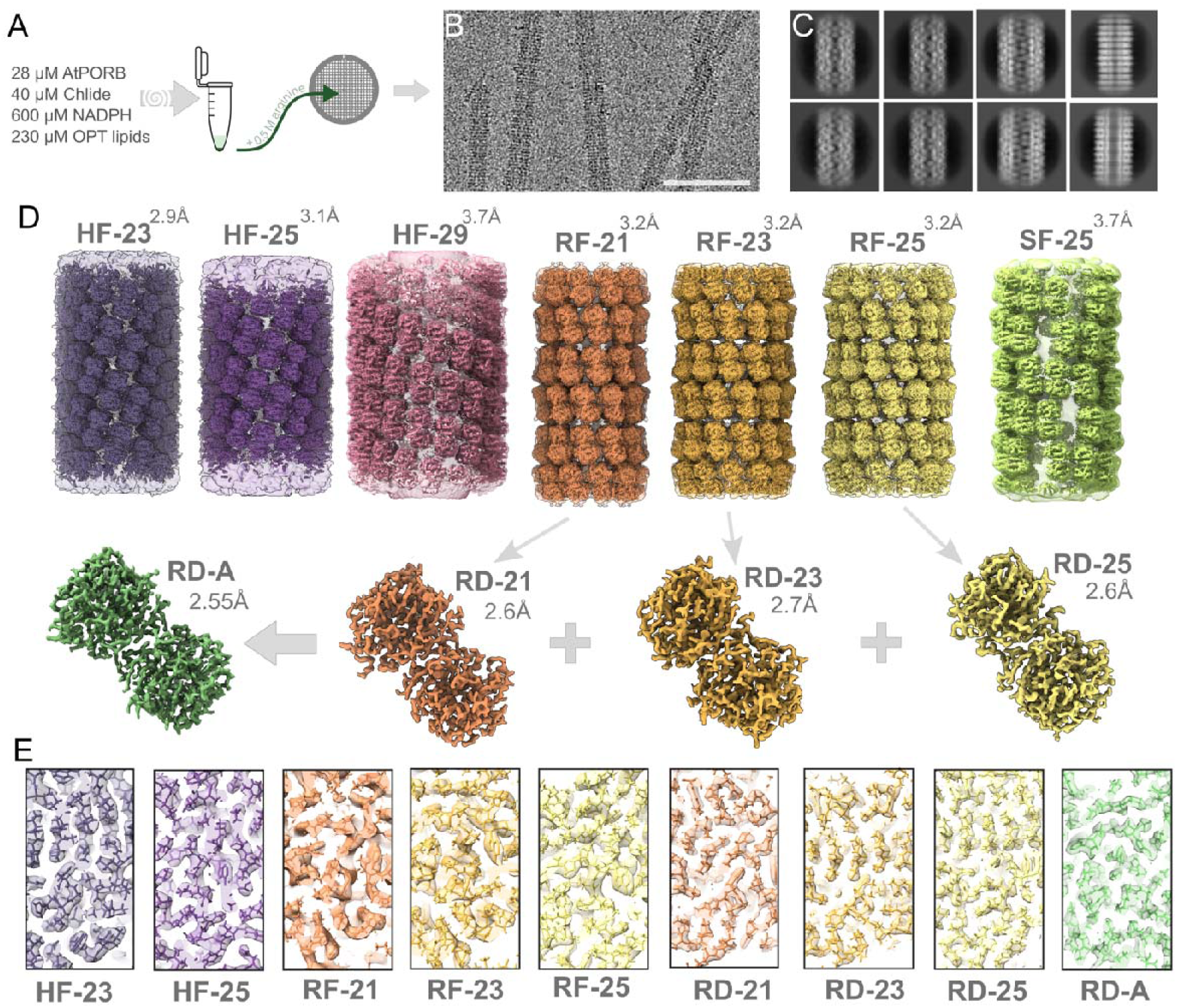
Cryo-EM reconstructions of Chlide:LPOR:NADPH oligomers. **A.** Simplified schematic of sample preparation. **B.** Representative cryo-EM micrograph; scale bar is 100 nm. **C.** Example 2D class averages. **D.** Cryo-EM maps of oligomeric assemblies from a single dataset, alongside dimer maps generated by sub-particle averaging; resolutions are indicated next to each map. Power spectra and the Fourier-shell correlation plots are presented in Extended Data Fig. 2. **E.** Cross-sections of each map superposed on the corresponding atomic model.

In total, we obtained eleven maps, enabling the construction of nine LPOR oligomer models; the HF-29 and SF-25 maps, however, lacked sufficient resolution for model building.

### LPOR forms strings of dimers that periodically deform membrane

To determine the mechanism by which LPOR can produce such a range of oligomeric structures, we investigated the lipid bilayer enclosed inside. Cross-sectional analysis of the assemblies revealed unexpected deformations of the bilayer. In top-down views, the helical assemblies displayed squarish and triangular cross-sections of the membrane for the 4-start and 3-start helices, respectively, which rotated along the filament’s long axis in accordance with the helical symmetry (Fig. 3A). In contrast, the ring-like assemblies appeared circular in top view but showed pronounced distortions inside cross-sections (Fig. 3A). The SF-assembly, on the other hand, was circular in a top view, but the center of the circle traced a helical path along the longitudinal axis.

**Figure 3.**
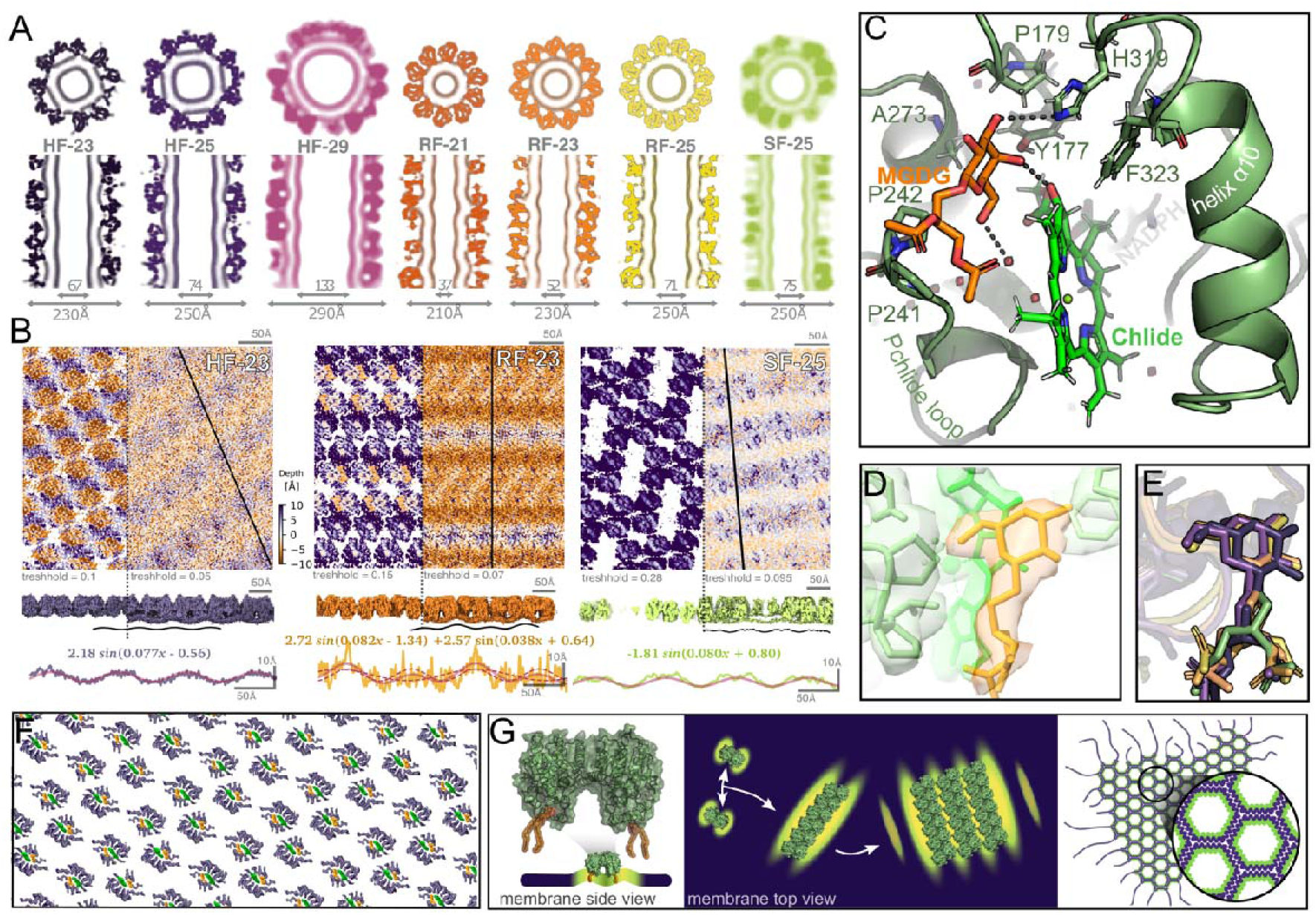
Strings of LPOR dimers induce periodic deformations of the bilayer enclosed within the filaments. **A.** Top-down and side cross-sections of cryo-EM maps, with filament and inner channel dimensions indicated. **B.** Planar projections of HF-23, RF-23 and SF-25 maps viewed from inside the filaments, with thresholds set to visualize either the monomer array or the outer leaflet of the membrane. The black line indicates the region used for profile analysis (10 nm window), with the experimental profile (violet, yellow or green), fitted curve (continuous red), individual components (dashed red), and fitting equation shown. **CD.** Detailed view on the MGDG binding site (C) and a model superimposed on the density map RD-A (D). **E.** Comparison of all MGDG molecules in all RD and all HF models. Colors correspond to panel A and Fig. 2D. **F.** Planar projection of HF-23 assembly, viewed from inside the filament, highlighting the array of MGDG molecules (yellow) and the pigment (green). Only helix α10 and Pchlide loop are visible. **G.** Schematic model of LPOR interaction with the membrane. Each dimer interacts with two MGDG molecules, inducing local membrane deformations (shape based on equation on panel B) that promote further oligomerization. These periodic deformations are likely essential for stabilizing the ultrastructure of PLB.

To better visualize membrane curvature, cryo-EM maps were projected onto planar representations using the unroll function in ChimeraX (Fig. 3B). This geometric transformation preserves the original density values and enables comparative analysis of surface features along the filament axis. In SF- and HF-type assemblies, all subunits are embedded at similar depths within the membrane, resulting in a regular, wave-like deformation of the outer leaflet that propagates perpendicular to the axis of the dimer string (Fig. 3BF). This pattern could be approximated by a single sine wave (R^2^=0.78 and 0.45 for HF-23 and SF-25, respectively). Interestingly, the membrane deformation was preserved within the SF-type assembly even in regions where the string of dimers was discontinuous, following the same single–sine-wave pattern (Fig. 3B). In RF-type assemblies, by contrast, the strings of dimers are tilted relative to the filament axis, which creates a zigzag-like pattern (Fig. 3B). As a result, the membrane surface is distorted in a more complex manner, requiring at least two sine waves to describe its curvature (R^2^=0.35): one with a period similar to that in helical assemblies, driven by the dimer length, and another with approximately twice the period, resulting from the zigzag orientation of the dimer strings.

Such a coordinated deformation of the membrane is possible because each LPOR dimer interacts with two MGDG molecules constraining it at defined angles and positions. The density of the head of the lipid is well resolved in RD-A map (Fig. 3D), but it seems that the orientation of MGDG is highly similar within all the determined assemblies (Fig. 3E). The lipid headgroup interacts primarily with the pigment and a water molecule, forming only a single hydrogen bond with the enzyme itself (residue H319) (Fig. 3C). This suggest that the interaction with the lipid, and therefore membrane remodeling, is possible only after the pigment binding.

Altogether, these data indicate that the underlying mechanism of membrane remodeling by LPOR is the induction of a deformation of the membrane caused by the dimers (Fig. 3G). A high MGDG content naturally induces a propensity for membrane curvature, but it is the scaffolding of strings of LPOR dimers, either continuous or segmented, that impose a coherent and directional deformation, probably inducing some shifts in MGDG distribution, what further stabilizes the periodic curvature. We therefore hypothesize that during PLB formation, strings of LPOR dimers induce and stabilize some specific membrane deformations that can propagate at a certain angle across a continuous network of curvature, stabilizing it and ultimately driving the formation of the cubic membrane phase characteristic of PLB architecture. To better understand the mechanisms underlying the curvature origin, we investigated the molecular architecture of the string of dimers.

### At least three different interactions between strings of dimers are possible

Remarkably, the dimer conformation is essentially identical across all assemblies we determined, including our previously determined Pchlide:LPOR:NADPH structure^9^ (Extended Data Fig. 3AD). By contrast, two dimers can interact at least in two distinct ways to form higher-order oligomers, which can be approximated as a square with one LPOR subunit at each corner (Fig. 4A). In RF-type assemblies, the square is nearly planar, whereas in HF-type assemblies the dimers are tilted, although the conformation of the interacting residues is almost the same as in RF-assemblies, except for the angle of interaction (Fig. 4B). This tilt can be represented by bending the square along one diagonal by an angle α: for RF-type assemblies α approaches 180°, while for HF-type models it is approximately 158° (Fig. 4C). We further define a second angle, β, which specifies the angle between adjacent squares along the string, forming a string. Across all models, β is consistently around 143° (Fig. 4C). This simple geometric model accounts for both helical and stacked-ring architectures (Fig. 4D).

**Figure 4.**
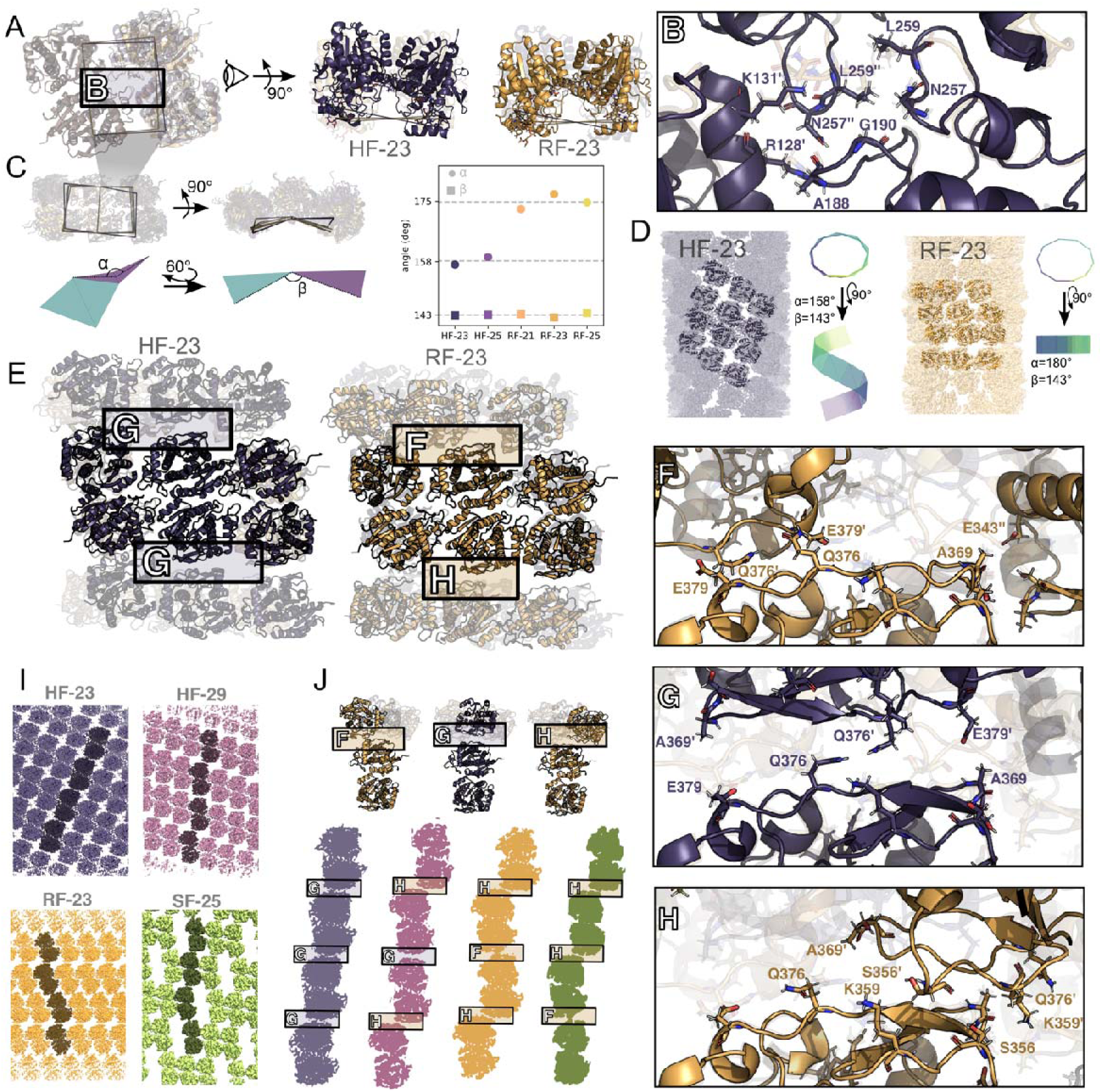
LPOR oligomers adopt architectures resembling sliding strings of squares. **A.** Superposition of two adjacent dimers from HF-23 and RF-23. **B.** Close-up of the HF-23 interface, with RF-23 shown semi-transparent for comparison. **C.** Geometric model representing the dimer string as a chain of squares bent along a diagonal; the two angles, α and β, are indicated, with their values calculated for each assembly. **D.** Twelve-mer models of HF-23 and RF-23 overlaid on their respective cryo-EM densities, alongside the idealized square-chain assemblies. Visualizations of string of squares on panels CD were generated by a simple mathematical model using the input α/β values. **E.** Overlay of HF-23 and RF-23 12-mers, highlighting three distinct oligomerization interfaces. **FGH.** Detailed views of the three interfaces marked in panel E. In all panels, the bottom subunit is shown in the same orientation to facilitate comparison of the relative position and shift of the upper subunit. **I.** Planar projections of selected maps, viewed from outside the filaments, with four dimers from adjacent strings highlighted in dark color. The interface types in each assembly are illustrated in vertically aligned four-dimer fragments of the corresponding maps. **J.** Superposition of all three interfaces, showing a central dimer interacting with a neighbor from an adjacent string.

To stabilize the oligomer, however, neighboring strings of dimers must interact. We identified three distinct interface conformations (Fig. 4E). In HF-23 and HF-25, only one symmetric interface is present, driven primarily by a mirror-symmetric interaction of Q376 residues (Fig. 4G). In RF assemblies, adjacent strings can associate through two distinct interfaces, depending on how the subunits are shifted relative to the mirror-symmetric arrangement. A leftward shift generates a looser contact involving K359:S356′ and Q376:A369′ (Fig. 4HJ), while a rightward shift produces a tighter interface involving Q376:E379′ and A369:E343″ from the neighboring dimer strand (Fig. 4FJ).

Interestingly, the same three different oligomerization interfaces can be found in HF-29 and SF-25 assemblies but arranged in different patterns. Unlike in HF-23 and HF-25, the mirror-symmetric interface (Fig. 4G) is present only between every other string of dimers, while the other strings are leftward shifted. In the SF-25 assembly, adjacent fragments of the dimer string are arranged in a repeating left–right–rightward shift pattern.

Comparison of these interfaces suggests that the dimer strings can slide relative to one another (Fig. 4J), as the polar and charged residues are arranged to permit multiple alternative contacts. Such variability in interaction patterns can give rise to a broad spectrum of combinations, resulting in distinct oligomeric architectures. These states are likely capable of dynamic interconversion facilitated by the inherent flexibility of the sliding interfaces.

### Pigment binding and the mechanism of the photoreaction

By combining all RD-type maps, we achieved a 2.55 Å resolution reconstruction (RD-A) of the LPOR dimer in complex with Chlide and NADPH, uncovering several unexpected insights into pigment binding and the reaction mechanism.

First, at this improved resolution, the Chlide ligand is now unambiguously resolved in the same orientation as in our earlier lower-resolution Pchlide:LPOR:NADPH map, for which the accuracy of the pigment orientation was verified with QM/MM modeling^9,27^ (Fig. 5A). Surprisingly, the surrounding binding-site residues adopt an identical conformation, despite now capturing the product rather than the substrate (Extended data Fig. 3). This implies that LPOR alone cannot easily distinguish between Pchlide and Chlide, indicating the existence of an additional mechanism for discriminating pre- and post-reaction states.

**Figure 5.**
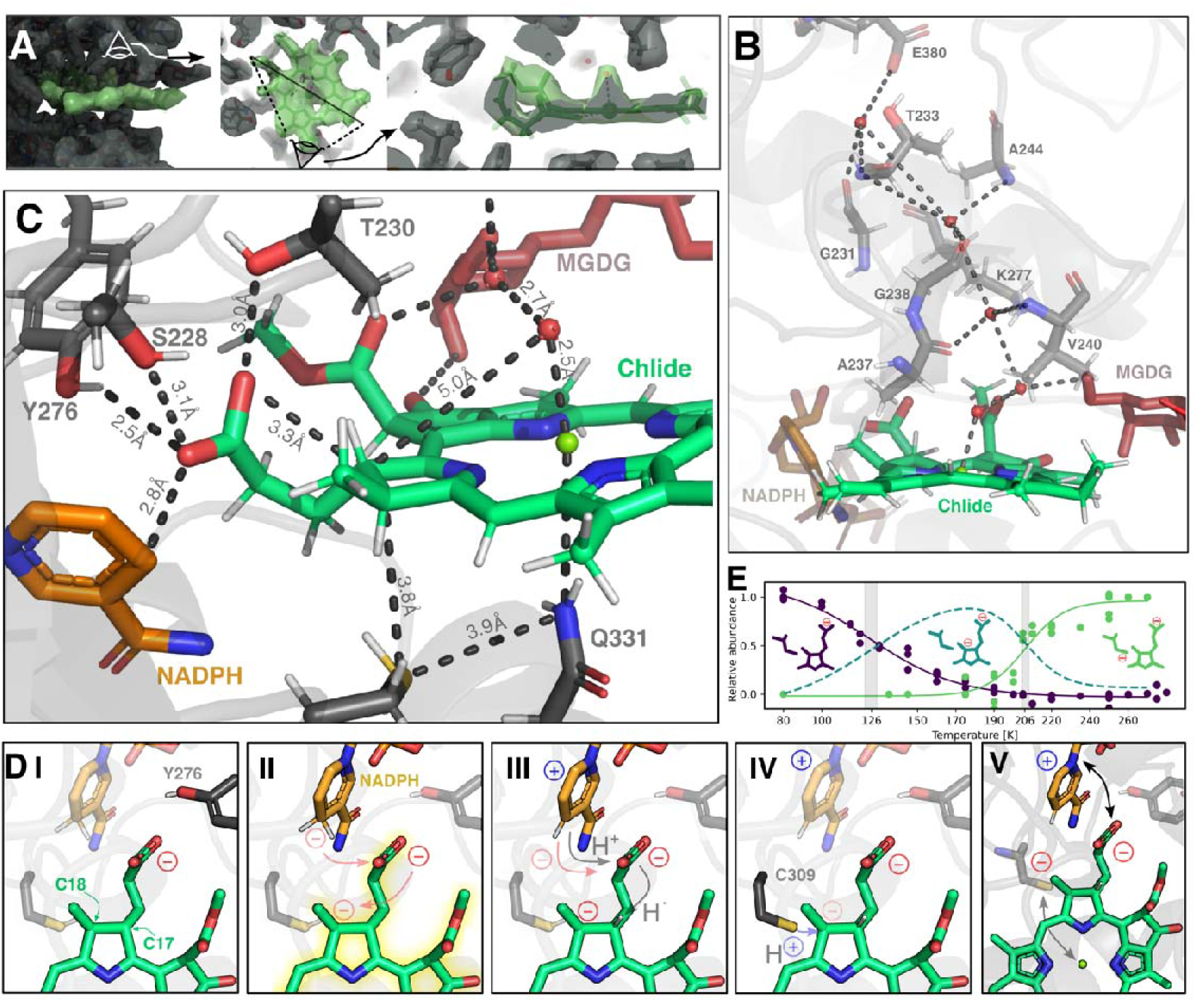
High-resolution RD-A map reveals a water channel and indicates a propionate-mediated hydride-transfer mechanism. **A.** Cryo-EM density of Chlide (green) in the RF-A reconstruction. A black line in a middle panel shows the cross-section shown on the next panel. **B.** Continuous chain of water molecules linking the pigment’s magnesium ion to bulk solvent via the Pchlide loop. **C.** Detailed view of the LPOR active site. **D.** Proposed reaction mechanism: I. Propionate at C17 is deprotonated in the ground state and awaits photon absorption. II. Upon photoexcitation, the electron density is redistributed, which triggers concerted electron transfers: from the propionate to C18, and from NADPH to the propionate, yielding an anion-radical intermediate. III. A proton from NADPH transiently neutralizes the carboxyl group, followed by a concerted transfer of a hydrogen radical from the propionate to C17 and an electron from NADPH to the propionate. IV. The negative charge at C18 is neutralized by a proton donation from the -SH group of C309, completing Pchlide reduction. V. The resulting thiolate coordinates the magnesium ion, inducing conformational change, while the propionate carbonyl engages NADP⁺ to promote further structural rearrangements. **E.** Relative abundances of substrate, intermediate, and product as a function of temperature, derived from the fluorescence data in Fig. 1A.

Second, we identified a water channel linking the pigment’s magnesium ion to bulk solvent through the Pchlide loop (Fig. 5B, Extended Data Fig. 4). We found five water molecules out of which one coordinates the pigment while interacting with backbone oxygen of A237. Another one interacts with both MGDG and pigment, while the rest of the water molecules are stabilized by numerous residues of the Pchlide loop.

However, the most striking finding was orientation of the propionate group at C17, which projects out of the tetrapyrrole plane to arch across the opposite face of the macrocycle (Fig. 5C). The carboxyl group is stabilized by Y276, S228, and T230, ideally positioned for hydride transfer from NADPH. We therefore propose a novel mechanism that begins with the propionate group already in its deprotonated state within the active site (Fig. 5DI, Extended Data Fig. 5). Then, as a result of photoexcitation, the electron density is redistributed across the tetrapyrrole, which induces a concerted transfer of electrons: from the propionate to C18 and from NADPH to the propionate (Fig. 5DII).). In this step, an anion-radical intermediate is formed. Subsequently, a proton from NADPH transiently neutralizes the carboxyl group of propionate, which is followed by a transfer of a hydrogen radical from the carboxyl to C17 and an electron migration from NADP⁺ to the propionate, that most likely occur again in a concerted fashion (Fig. 5DIII). Then, the resulting negative charge at C18 promotes a proton transfer from C309, what completes the reduction of Pchlide (Fig. 5DIV). The resulting thiolate group at C309 then competes with a water molecule on the opposite face for magnesium coordination, driving pigment displacement, conformational change, and product release (Fig. 5DV). Concurrently, the propionate carbonyl at C18 may interact with the oxidized nicotinamide ring of NADP+, triggering further structural rearrangements. In the enzyme’s open state, C309 could be reduced by either a solvent water, Y276 or Y306.

In this model, photoexcitation drives intramolecular charge migration culminating in thiolate formation, which then signals product release. This explains how LPOR discriminates the post-reaction Chlide complex from the merely bound state: only immediately after catalysis C309 bears a negative charge, what triggers conformational change leading to dinucleotide release.

Moreover, by monitoring Pchlide and Chlide fluorescence during the temperature-resolved reaction (Fig. 1A, 5E), we pinpointed the activation temperatures for each step. Hydride transfer becomes efficient at ∼126 K, while thiolate formation requires ∼206 K (Fig. 5E). Thus, illuminating native LPOR complexes within PLB at ∼170 K traps the reaction intermediate, providing an optimal condition for experimentally testing our proposed mechanism.

## Discussion

### [NADPH]/[NADP^+^] ratio drives the formation and disassembly of PLB

Our data indicate that PLB disassembly is delayed by the formation of Chlide:LPOR:NADPH complexes, driven by the high NADPH concentration present in dark-incubated etioplasts and immediately after illumination. Although seedlings would be expected to initiate assembly of photosynthetic machinery as rapidly as possible, this transient stabilization may act as a safeguard against dismantling PLBs after brief, incidental light exposure or could play a photoprotective role. Together with our previous observation that Pchlide:LPOR:NADP⁺ complexes promote the formation of a cubic-like phase in vitro^8^, these results highlight the central role of changes in the NADPH/NADP⁺ concentration ratio in both the development and disassembly of PLBs (Fig. 1D). We propose the following sequence: early in PLB biogenesis, high [NADP⁺] favors an irregular cubic phase; as [NADPH] rises, assembly of the active LPOR complex proceeds and the paracrystalline PLB ultrastructure emerges. Following photoreduction of Pchlide, thiolate formation in the binding pocket triggers NADP⁺ release. Then NADPH rebinds in the LPOR binding pocket, stabilizing an LPOR oligomers with Chlide and transiently delaying PLB disassembly. Dinucleotide displacement is likely accompanied by subunit rearrangements within oligomers, which could explain the partial deformation of the PLB ultrastructure observed after illumination^28^. The eventual release of Chlide, known as the Shibata shift, likely requires a decline in NADPH and/or the action of additional regulatory proteins. Consistent with this view, the [NADPH]/[NADP⁺] ratio was shown to decrease from ∼4 in dark-incubated etioplasts to ∼0.6 after 30 min of illumination, but a significant change was detected already after 2 min^29^. This suggests that some additional light-activated processes consuming NADPH drive the redox transition that leads the assembly of photosynthetic machinery.

### LPOR can play regulatory role both in developing and mature leaves

The results presented here show that LPOR:NADPH forms stable complexes with both Pchlide and Chlide, but only the Chlide-bound complexes remain stable upon illumination, effectively trapping this chlorophyll intermediate. This links Chl synthesis to photosynthetic activity: elevated NADPH concentration generated by photosystems can pause the flux at the terminal step. Recent work further shows that plant LPOR isoforms differ in their affinities for NADPH and Chlide^30^, implying that plants can tune the rate of Chl synthesis by modulating the expression of LPOR isoforms with distinct biochemical properties. This regulatory role of LPOR likely integrates with the broader plastid redox network, including ferredoxin and thioredoxin systems^31^, out of which the latter is known to modulate enzymes of the Chl biosynthetic pathway^32,33^.

Moreover, because LPOR oligomers are essentially the same whether they contain Pchlide or Chlide, it is plausible that both pigment-bound states can engage FLU, the principal negative regulator of Chl biosynthesis that inhibits the first enzyme in the pathway^34–36^. To date, FLU has been shown to interact primarily with Pchlide:LPOR:NADPH complexes^37^; but our findings suggest that the FLU:LPOR feedback loop may govern not only Pchlide abundance in etiolated seedlings but also the daytime rate of Chl synthesis in mature leaves.

### Is periodic deformation a universal mechanism of remodeling of photosynthetic membranes?

Thylakoid-lipid extracts assemble into lamellar bilayers in aqueous buffer^38^, yet their high MGDG content confers a pronounced negative-curvature propensity^39,40^. The structures of the LPOR assemblies described here indicate that the enzyme can harness this predisposition by imposing small, periodic deformations, steering MGDG-rich membranes into distinct, self-stabilizing morphologies. We therefore hypothesize that periodic, protein-driven deformation is a general mechanism of photosynthetic membrane remodeling. Consistent with this view, VIPP1 oligomerizes into interconnected dome-shaped complexes and filamentous assemblies that confine an undulated membrane^41,42^. We further speculate that CURT1A, which is thought to stabilize grana margins ^43^, may act similarly; notably, its overexpression tightens PLB ultrastructure^44^, suggesting that, together with LPOR, it could impose additional periodic deformations that modulate the cubic phase. More broadly, this hypothesis opens the door to designing de novo proteins that reshape membranes into defined morphologies by exploiting the same physical principle.

### A Novel Reaction Mechanism Consistent with Previous Observations

The structural data presented in this paper suggest that the underlying role of photoexcitation is charge migration from the propionate group at C17 to C309, which is accompanied by the proton and electron transfer from NADPH. Determining the exact order of events will require further study and QM/MM calculations; nevertheless, although this reaction mechanism has not been examined directly before, it is consistent with previous work supporting a charge-transfer scenario^45^ and with the experiment pinpointing the C17 as the NADPH-reduced carbon^46^. Notably, the stepwise nature of the reaction mechanism was proposed several decades ago, featuring a non-fluorescent intermediate stable at 175 K^47^, which agrees with the data presented here (Fig. 1A, 5E). Similar results were observed for cyanobacterial LPOR variants in vitro, although the activation temperature for intermediate formation was higher than reported here^16^. Notably, both hydride and proton transfers were shown to be faster in eukaryotic LPOR isoforms than in cyanobacterial ones^48^, which may explain this discrepancy and suggests differences in the orientation of the propionate group between these LPOR types.

The central role of the propionate in the LPOR reaction mechanism is surprising; nevertheless, its importance for the activity has been demonstrated before, using chemically modified pigments^49^. This mechanism also clarifies the significance of Y276, long suspected to be a proton donor. The residue, together with S228 and T230, orient the propionate carboxylate in the active site. Accordingly, mutation of either residue diminishes, but not abolish enzyme activity^9,15,50^.

Altogether, our findings shed new light on the regulatory role of LPOR in mature leaves, the enigmatic photoreaction mechanism, and membrane remodeling in developing chloroplasts, offering design principles for engineering plastid membranes.

## Methods

All procedures that involved the seedlings or LPOR reaction mixtures at room temperature described below were carried out in darkness or under diffuse, non-actinic green safelight to avoid initiating LPOR photochemistry.

### Etiolated seedlings and their low-temperature measurements

Etiolated seedlings of Triticum aestivum were grown in the dark for 7 days at room temperature. Seedlings of similar size were selected and, after removing 0.5 cm from the tip, the subsequent 2 cm segment was used for the experiments.

Temperature-series fluorescence: Fluorescence spectra were recorded in a liquid-nitrogen bath cryostat (OptistatDN, Oxford Instruments) coupled to an Ocean Optics QE65000 spectrometer and equipped with a 600-nm long-pass filter (Edmund Optics). Excitation was provided by an Leica Schott KL 1500 LCD lamp with a 456 ± 5 nm band-pass filter (Carl Zeiss Jena). Spectra were acquired at discrete temperatures from 80 K to 280 K. For each target temperature after the initial 80 K measurement (without prior illumination), the sample was illuminated for 1 min with saturating white LED light (> 6000 μmol photons m⁻² s⁻¹) and then dark-incubated for 1 min at the target temperature. At least two fluorescence spectra were collected at each temperature for each seedling. In total, the temperature-series experiment included 3 seedlings and 64 spectra.

Time-course fluorescence: the measurements were performed on a PerkinElmer LS-50B fluorimeter equipped with a liquid-nitrogen-cooled sample holder. The seedlings were placed in glass tubes and then illuminated with white light (5 μmol photons m⁻² s⁻¹) for specified durations (1 s–40 min) and immediately frozen in liquid nitrogen. Spectra were recorded at 77 K from 600 to 790 nm with a scanning speed of 400–500 nm min⁻¹, a data interval of 0.5 nm, and an excitation wavelength of 440 nm. Excitation and emission slit widths were 7–10 nm. For each seedling, spectra were recorded in triplicate with 120° rotations between acquisitions. Data analysis and Gaussian fitting were performed using Python scripts employing Matplotlib and NumPy. All spectra were normalized to their maxima. In total, the time-course experiment included 48 seedlings and 144 spectra.

### Pigments, recombinant protein and in vitro experiments

Pchlide was purified from etiolated wheat seedlings according to the protocol published by the authors^11,51^. Chlide was produced from spinach chlorophyll using recombinant chlorophyllase as described by one of the authors^52^.

Recombinant AtPORB from Arabidopsis thaliana (UniProt: P21218) was expressed in E. coli and purified via His-tag affinity chromatography as described previously^30,53^. For in-vitro LPOR activity measurements, reaction mixtures contained 37 mM sodium phosphate buffer (Na₂HPO, pH 7.1), 225 mM NaCl, 150 mM imidazole, 5 mM β-mercaptoethanol, and 25% (v/v) glycerol. Exact reagent concentrations are provided in the figure legends. “OPT lipids” denotes the lipid mixture optimized in our previous structural work^9^, comprising 50 mol% MGDG, 35 mol% DGDG, and 15 mol% PG (Avanti Polar Lipids).

Reaction mixtures were prepared by diluting the LPOR stock to 15 µM, adding NADPH (AppliChem), then adding the OPT lipid mix (MeOH stock) directly to the reaction using a micropipette, followed immediately by vortexing. Pigment, either Pchlide or Chlide, was added last, followed by vortexing. Samples were incubated for 20 min in darkness and transferred to glass tubes. Reactions were initiated with white light (5 μmol photons m⁻² s⁻¹) for specified durations (1 s–40 min) and stopped by immediate freezing in liquid nitrogen. Fluorescence measurements were performed on a PerkinElmer LS-50B as described above. Data analysis was performed using Python scripts employing Matplotlib and NumPy. Together, the kinetic experiment in vitro involved 84 samples/spectra.

Selected samples were negatively stained according to the previously published protocol^9^ and visualized with JEOL JEM2100 HT CRYO LaB6 electron microscope. At least three independent sample preparations were analyzed, and representative micrographs are presented.

### Preparation of cryo-EM sample

Sample composition was optimized by negative-stain TEM before preparing cryo-EM samples. An arginine solution (0.5 M arginine in phosphate buffer, pH 7.1, containing 225 mM NaCl and 5 mM β-mercaptoethanol) was added to the AtPORB stock to reach final concentrations of 150 mM arginine and 28 µM LPOR. NADPH was then added to 300 µM, followed by 230 µM OPT lipids; the mixture was vortexed immediately after each addition. Finally, 40 µM Chlide (in MeOH) was added with immediate vortexing; the final MeOH content was <6% (v/v). Duplicate samples were incubated for 7 min in darkness at room temperature and centrifuged at 21,300 g for 7 min at 4 °C. The supernatant was discarded and the pellet was immediately resuspended in 10 µL of the arginine solution. Resuspended pellets from two samples were combined, and 3.5 µL of the mixture were applied to Quantifoil R2/1, Cu 200-mesh grids. Grids were glow-discharged immediately before use (60 s, 8 mA) and vitrified on a Vitrobot Mark IV (Thermo Fisher Scientific) under the following conditions: 100% humidity, 277 K, blot time 6 s, wait time 0 s, blot force 5, drain time 0 s, one blotting cycle. After plunge-freezing in liquid ethane, grids were stored in liquid nitrogen until use.

### Data acquisition and processing

Cryo-EM data (18 466 movies, 40 frames each) was collected at the National Cryo-EM Centre SOLARIS in Kraków, Poland. Dataset was collected on a Titan Krios G3i microscope (Thermo Fisher Scientific) at an accelerating voltage of 300[kV, a magnification of ×105,000 and a pixel size of 0.86 Å using EPU 2.10.0.1941REL software. A K3 direct electron detector was used for data collection in a BioQuantum Imaging Filter (Gatan) set-up with a 20[eV slit width. The detector was operated in counting mode. Imaged areas were exposed to 40[e−/Å2. The defocus range applied was −2.1 µm to −0.9 µm with 0.3 µm steps. All datasets were processed using cryoSPARC v4.4.049–5353–56. Micrographs were motion-corrected using Patch Motion Correction, and the contrast transfer function (CTF) was determined using Patch CTF.

For particle picking, an initial set of ∼600 segments was manually selected to generate 2D class averages, which were then used to re-pick a subset of ∼2,000 micrographs with the Filament Tracer job. This provided 2D class averages that captured the high heterogeneity of filament geometry (42 references selected, filtered to 15 Å). A final reference-based Filament Tracer job was then performed on the entire dataset. Approximately 4 million segments were extracted with a box size of 640 × 640 pixels and an inter-box distance of 22 Å, and subsequently downsampled to 128 × 128 pixels for iterative 2D classification, until a clean set of 2.6 million segments was obtained. The corresponding 2D class averages were first manually sorted by filament diameter, and then, within each diameter group, the average power spectra of the segments belonging to each class were computed to further separate the data into distinct helical types with different symmetries, since several symmetries co-existed for the same diameter. Using a representative average power spectrum for each helix type, Helixplorer-1 was used to estimate the helical parameters (Extended Data Fig. 2), which allowed us to define two main populations: the Ring Filaments (RF), characterized by a large helical rise of ∼155 Å and high rotational symmetry (C10–C12), and the Helical Filaments (HF), which displayed smaller rises (6–25 Å) and lower rotational symmetries (up to C4). Candidate solutions for the helical parameters and symmetries determined in Helixplorer-1^54^ were subsequently validated by a Helical Refinement job^57^. In the first round of refinement, no additional dihedral (D) symmetry was imposed; this was only applied in a second step, once the presence of such symmetry had been confirmed. In all cases, the initial confirmation of helical symmetries was performed on downsampled particles (pixel size 1.72 Å). All structures, with the exception of the HF-29 filaments, exhibited dihedral symmetry, i.e. they are apolar. Once the symmetries had been validated, a final refinement with unbinned particles was carried out, yielding the final maps for HF-23 and HF-25, whereas the other structures required additional steps as detailed below.

For the HF-29 filaments, in the initial map, where the correct helical parameters had been applied, only two of the three strands were well resolved, while the third appeared blurred, giving the impression of two overlapping possible positions. To address this, a 3D classification without alignment, starting from the orientations obtained in the first refinement, was performed. The classification split the dataset into two equally populated classes that differed only by a 180° rotation perpendicular to the helix axis. This revealed that an ambiguity in polarity (in-plane angle assignment of the segments) was responsible for the poor quality of the initial refinement: two of the three strands were apolar and therefore aligned consistently, whereas the third strand disrupted this apolarity and became blurred because the segments could adopt either in-plane 180° rotated orientations. Since both 3D classes showed a well-resolved third strand, we used one as a reference for a final refinement against all particles, which yielded the final map.

For the Ring Filaments, the large helical rise exceeded the inter-box distance of the initially extracted segments, which introduced data redundancy in the first refinements. To address this, we performed a new template-based Filament Tracer job, using 2D projections of the initial HF-21, HF-23, and HF-25 structures, and extracted segments with the same box size as before but with an inter-box distance of 155 Å. This yielded 369,000 particles, which were iteratively 2D classified down to 72,000 segments corresponding to the three structures. To separate these, a Heterogeneous Refinement was carried out using the previous initial 3D maps as references, resulting in 22,000, 25,000, and 25,000 particles for HF-21, HF-23, and HF-25, respectively.

After an initial helical refinement, subsequent steps were conducted by treating the segments as single particles, with symmetries of D10, D11, and D12, since extraction at every helical rise allowed us to relax the helical symmetry constraint. A mask enclosing only the central double-ring of dimers was then applied for a Homogeneous Refinement, followed by per-particle defocus refinement and Local Refinement with updated CTF parameters.

To further improve resolution, we employed a symmetry-expansion and density-subtraction strategy, merging dimers from the three structures into an expanded dataset to correct for potential variability in dimer positioning. Specifically, after symmetry expansion (D10, D11, and D12, increasing dataset size by more than 20-fold), a mask was applied to subtract all densities except a single dimer, and local refinement was performed with a mask enclosing that dimer. To align the three resulting reconstructions of a single dimer from the RF maps, the Volume Alignment Tools job was used to bring their centers of gravity to a common origin while updating particle orientations into this common reference frame. Finally, the symmetry-expanded, subtracted, and aligned particles corresponding to the three Ring Filament types were merged for a Local Refinement, yielding the highest-resolution dimer map from this dataset (2.6 Å). All resolution estimates were based on gold-standard FSC values at the 0.143 cut-off.

## Acknowledgments

This work was supported by SONATA project (2019/35/D/NZ1/00295) granted by National Science Centre (NCN) to MG. The cryo-EM study was supported under the Polish Ministry of Education and Science project: ‘Support for research and development with the use of research infrastructure of the National Synchrotron Radiation Centre SOLARIS’ under contract number 1/SOL/2021/2. Data collection was performed under proposal nr. 221010. We also acknowledge the Polish high-performance computing infrastructure PLGrid (HPC Center: ACK Cyfronet AGH) for providing computer facilities and support within computational grant numbers: PLG/2023/016449, PLG/2024/017296 and PLG/2025/018450.

## Author contributions

The study was designed and directed by M.G.

L.F. provided Chlide, M.G. and J.K purified Pchlide.

A.G. and M.Z. performed low-temperature fluorescence measurements.

M.G. optimized and prepared sample of Chlide. Grids freezing and initial grid screening was performed by M.R., G.W. and M.G. Data collection was done by M.R. and G.W. Data processing was performed by A.D., L.F.E. and S.P.

Figures were prepared by M.G except Extended Data Fig. 2, which was prepared by L.F.E. The paper was written by M.G. with all authors discussing the results and refining and approving the final version.

## Data availability

The cryo-EM densities have been deposited in the Electron Microscopy Data Bank (EMDB) and the protein models have been deposited in the Protein Data Bank (PDB) with the following accession codes: RD-A: 9TL8 / EMD-56045; RD-21: 9TL6/ EMD-56043; RD-23: 9TL7 / EMD-56044; RD-25: 9TL9 / EMD-56046; RF-21: 9TLA / EMD-56047; RF-23: 9TLB / EMD-56048, RF-25: 9TLC / EMD-56049; HF-23: 9TLH / EMD-56055; HF-25: 9TLK / EMD-56056; HF-29: EMD-56051, SF-25: EMD-56050.

## Notes

### Competing Interest Statement

The authors have declared no competing interest.

