## Extended Data for "Structures of the LPOR–Chlide Complexes Imply the Basis of Membrane Remodeling and a Photocatalytic Mechanism"

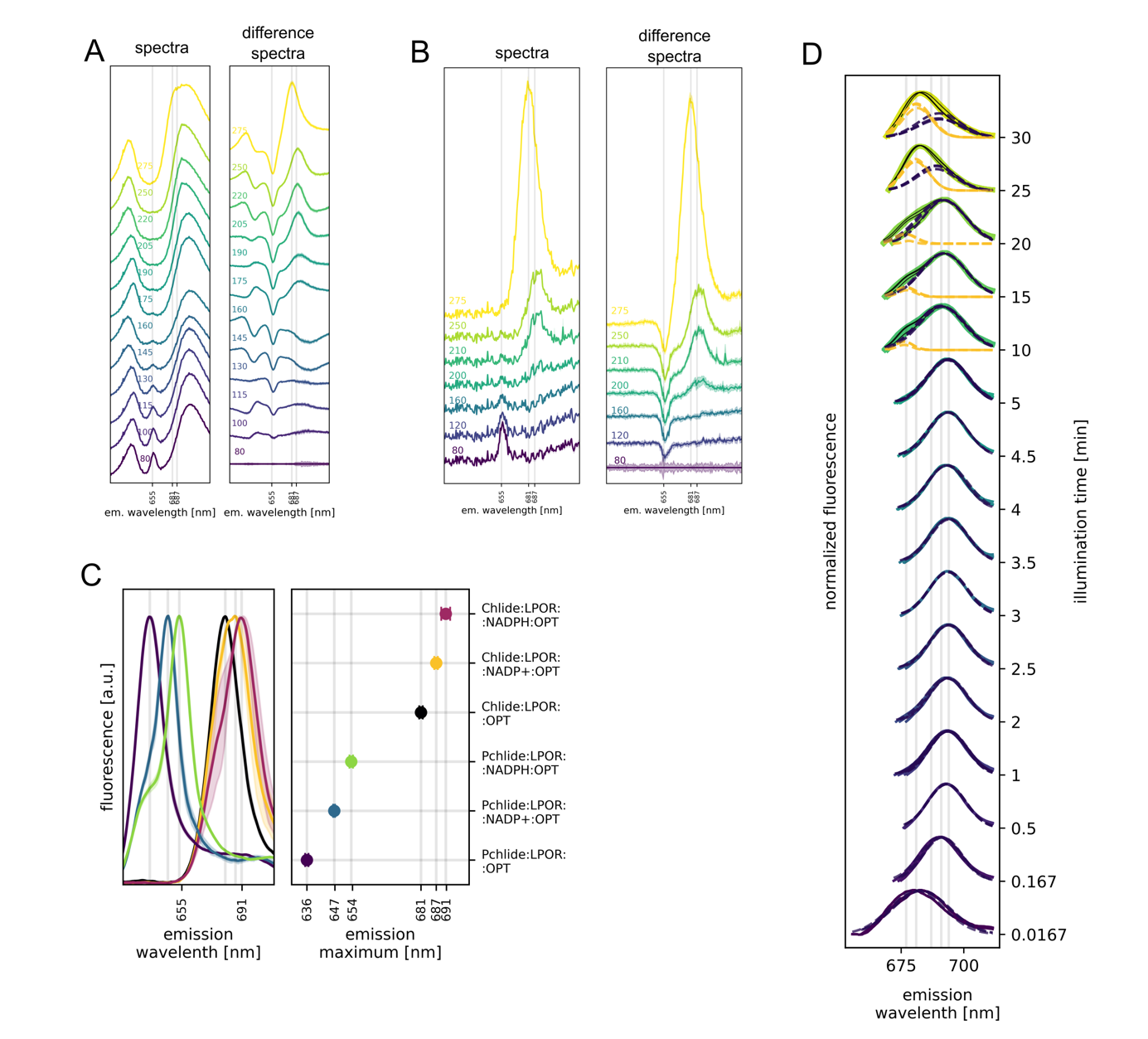

**Extended Data Figure 1. AB.** Fluorescence spectra of wheat seedlings illuminated over a temperature range. Shaded areas represent standard deviation between biological replicates. On A, peak at ~720nm originates form a light contamination. **C.** The fluorescence emission spectra of LPOR (15µM) in complex with NADPH/NADP+ (200 µM) and Pchlide/Chlide (5 µM), in the presence of OPT lipids (400 µM), with emission maxima corresponding to complexes of different compositions. **D.** Difference fluorescence spectra of wheat seedlings (black) illuminated for varying time intervals and measured at 77 K with Gausian deconvolution (yellow dashed line – components (if present), violet dashed line – sum of components). Shaded area represents standard deviation between biological replicates.

A

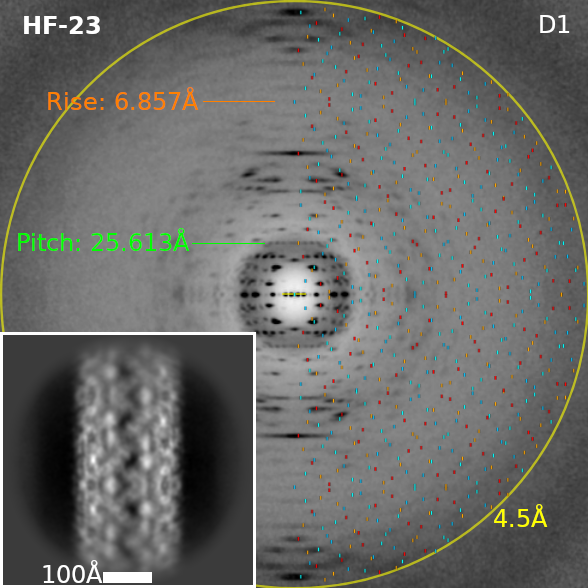

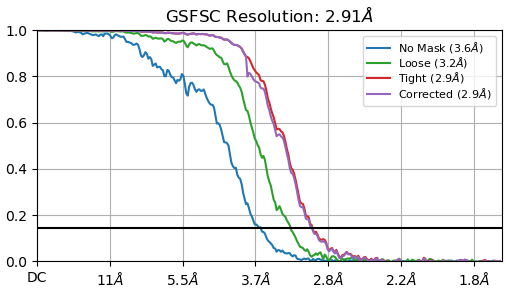

B

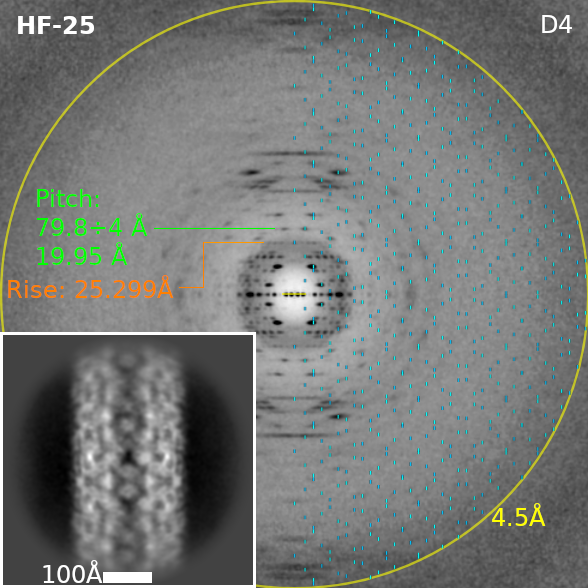

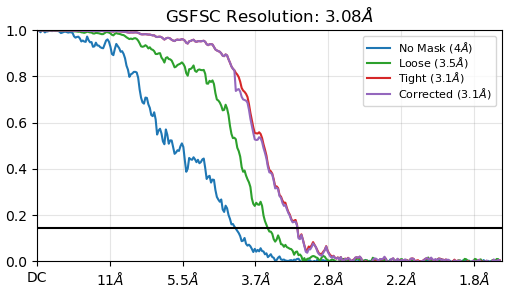

C

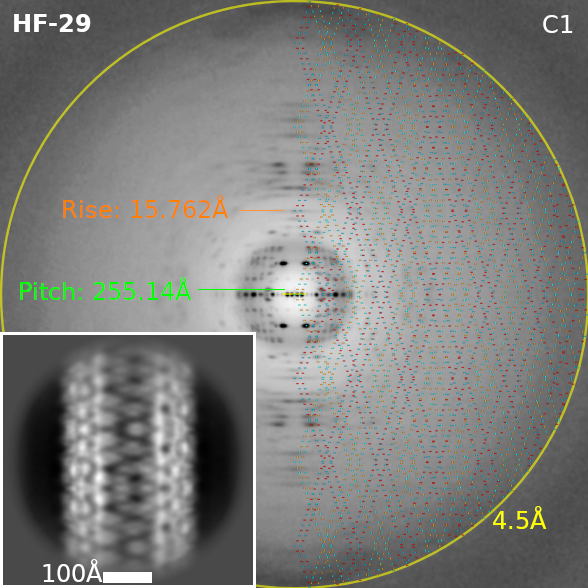

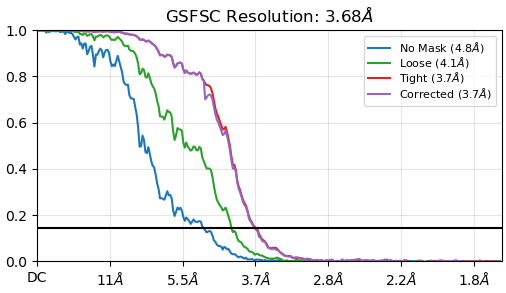

D

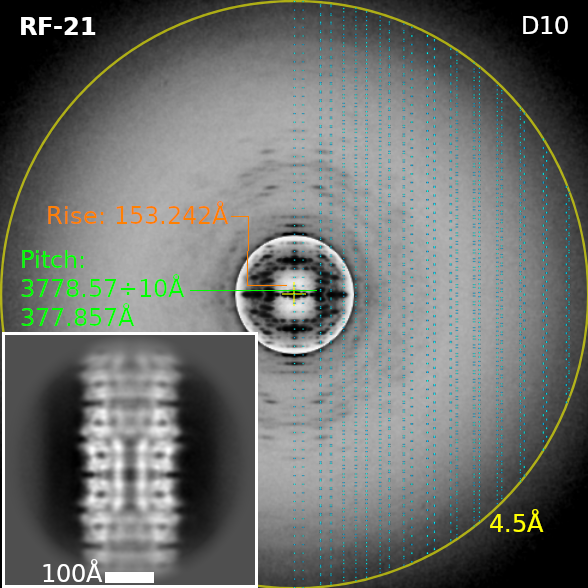

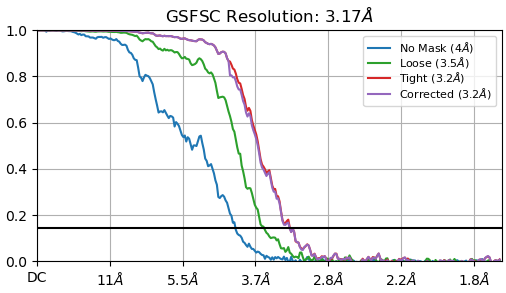

E

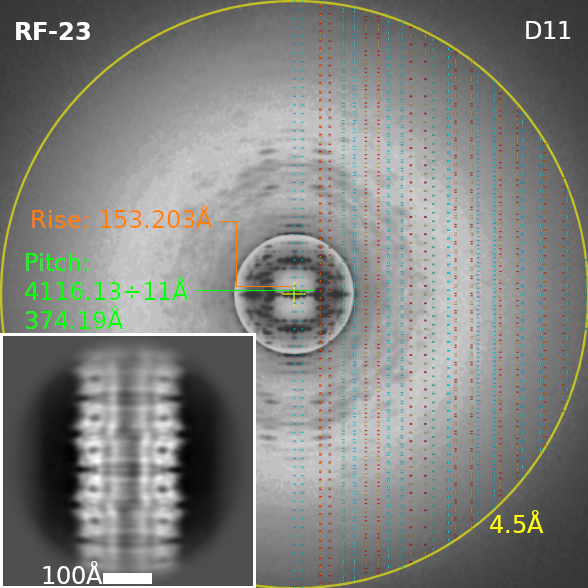

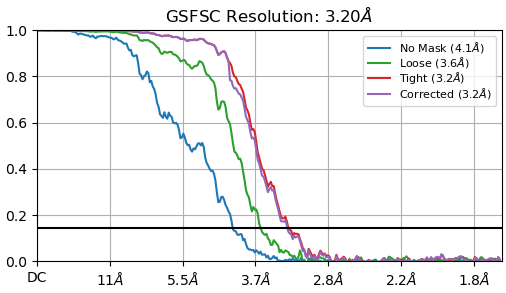

F

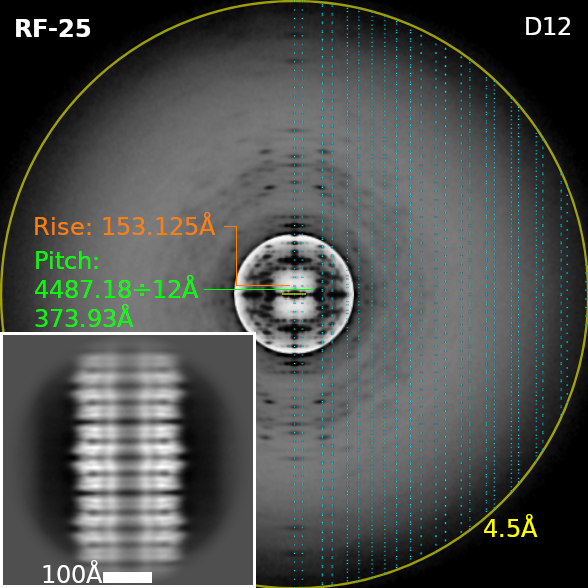

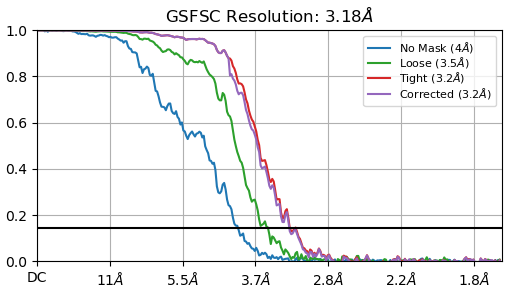

G

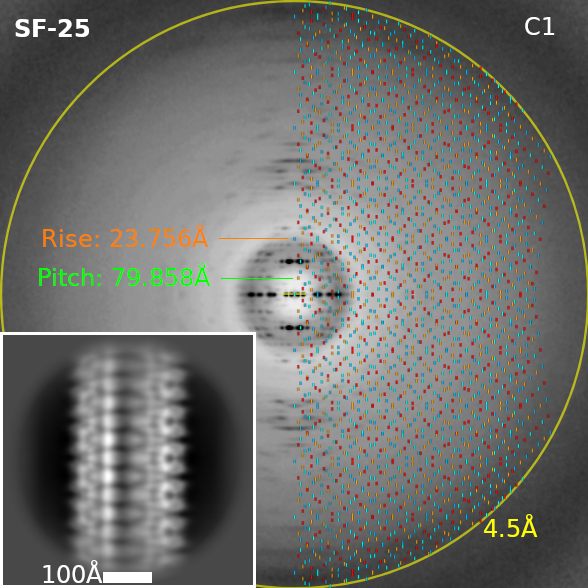

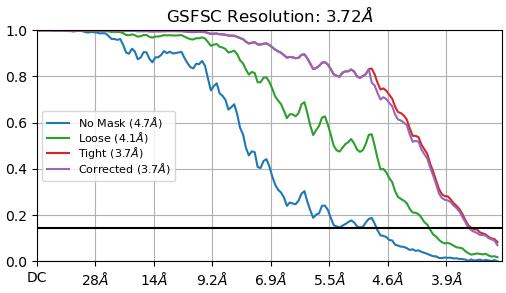

H

**RD-21**

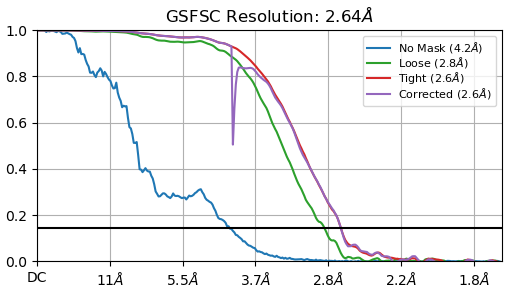

I

**RD-23**

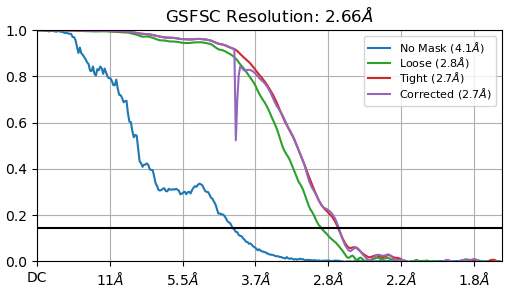

J

**RD-25**

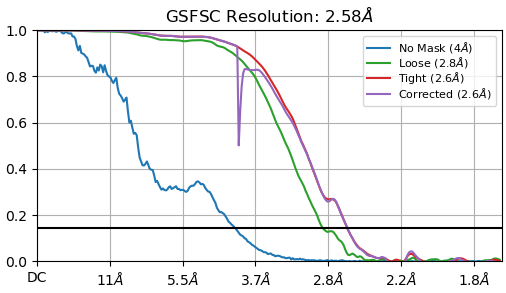

K

**RD-A**

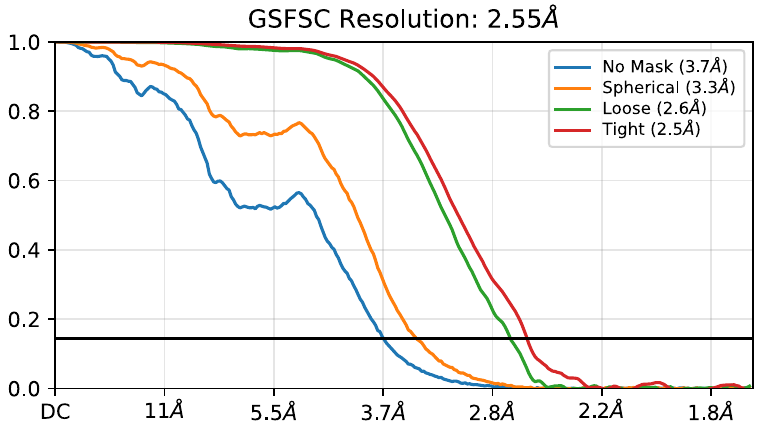

**Extended Data Figure 2.** Helixplorer^57^ symmetry determination plots and resolution estimation by gold-standard Fourier-shell correlation (GSFSC) of HF-23 (**A**), HF-25 (**B**), HF-29 (**C**), RF-21 (**D**), RF-23 (**E**), RF-25 (**F**) and SF-25 (**G**) structures as well as resolution estimation by gold-standard Fourier-shell correlation (GSFSC) of RD-21 (**H**), RD-23 (**I**), RD-25 (**J**) and RD-A (**K**). On the left: average power spectra (PS) in inverted gray-scale (stronger spots are black) for each type of helical assembly overlaid with the theoretical positions of Bessel function maxima. These maxima positions depend on the helical symmetry parameters used (pitch and axial rise). The parity of Bessel function orders are conditioned by the parity of additional cyclic/dihedral symmetry (Cn/Dn) that might also be present. Small blue and red dots on the right side of the PS indicate the expected positions of even and odd Bessel orders maxima, respectively. Insets show representative 2D class averages for each type of helical assembly. The intensity of spots in the PS depend both on the size, shape and orientation of the asymmetric unit and on electron microscope contrast transfer function (CTF). The CTF strongly attenuates low spatial frequencies near the PS center which explain why the pitch spots in (C) and the axial rise spots in (D,E,F) are not visible in the PS. On the right: Fourier shell correlation curves for the reconstructions presented in Figure 2 with different kinds of masks applied.

^57.^	Estrozi, L.F., Desfosses, A., Schoehn, G., 2018. Helixplorer: Online Indexation of Fibers And Helical Structures, <http://rico.ibs.fr/helixplorer/>

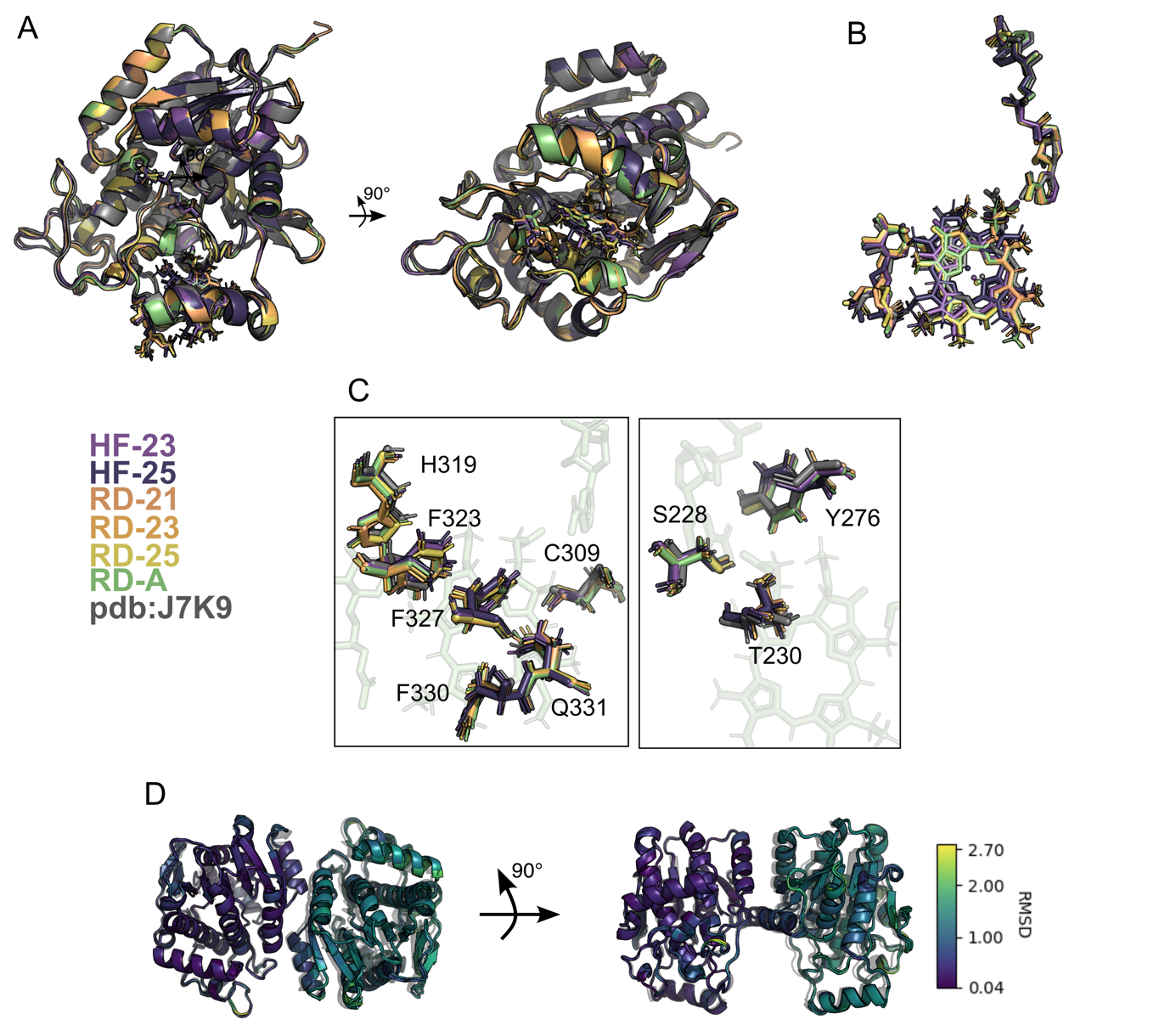

**Extended Data Figure 3. LPOR dimers in all assemblies are essentially identical.** **A.** Comparison of subunits from all assemblies, including the subunit of Pchlide:LPOR:NADPH complexs from pdb:7JK9**. B.** Overlay of NADPH, Pchlide and MGDG in LPOR subunit form all assemblies.

**C.** Conformation of selected residues involved in Pchlide/Chlide binding in subunits from all assemblies, including the subunit of Pchlide:LPOR:NADPH complexs from pdb:7JK9. **D.** Comparison of the conformation of LPOR dimers form all assemblies aligned to the dimer from PDB:7JK9 (in black semitransparent). The RMSD for each residue is shown in a color scale.

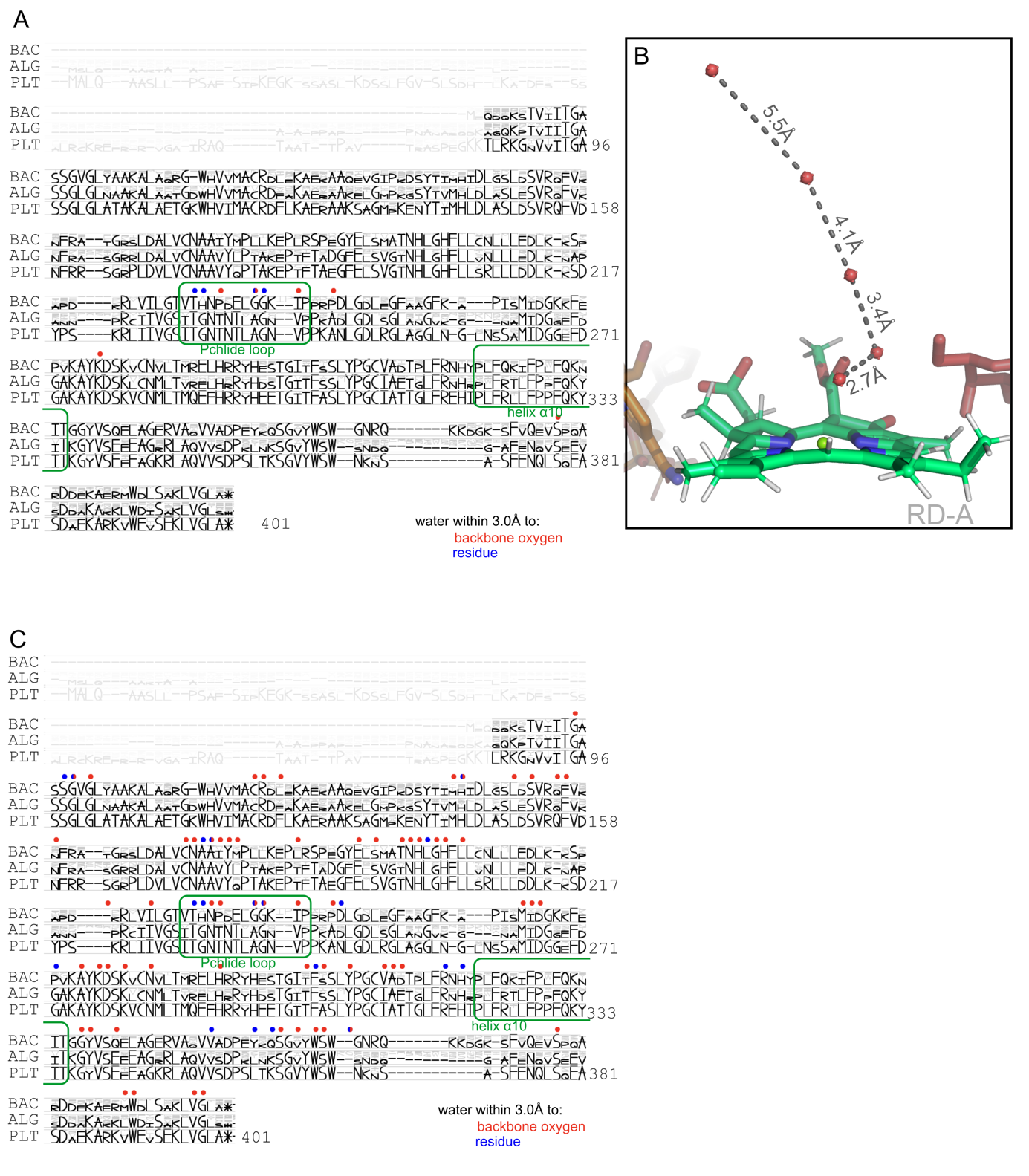

**Extended Data Figure 4. Residues involved in water binding.** Sequences 346 of cyanobacterial (BAC), 19 algal (ALG), and 190 plant (PLT) aligned LPOR sequences. The size of each letter reflects the abondance of the given residue at a given position. **A.** Residues interacting with the water channel connecting the pigment with bulk solvent. **B.** The distances between the water molecules in the channel. **C.** The residues interacting with all water molecules present in RD-A model.

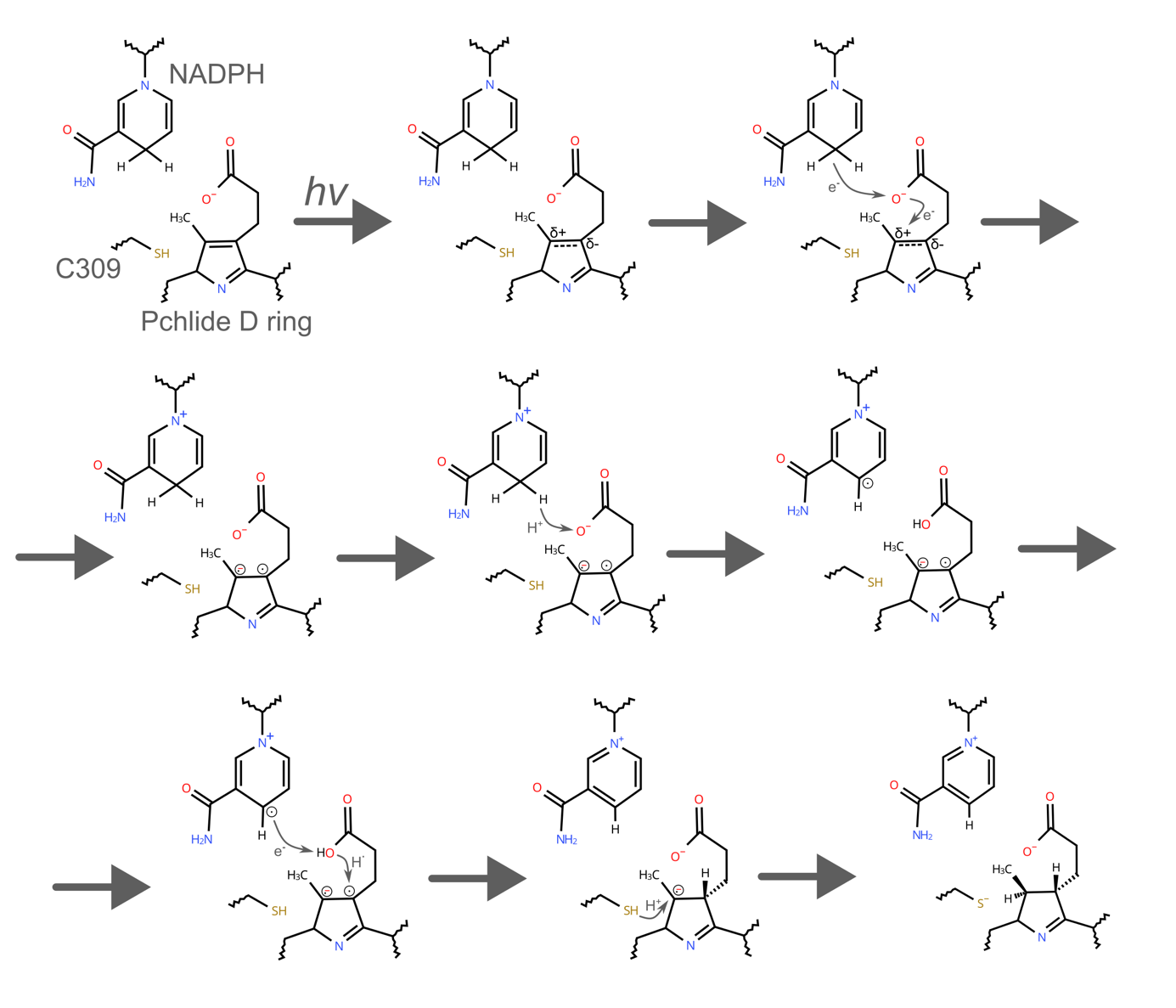

**Extended Data Figure 5. A novel reaction mechanism of LPOR.**

**Supplementary video 1.** Overview of the maps determined in this study.

| **Extended Data Table 1. RD-A cryo-EM data collection and refinement statistics.** | |
| --- | --- |
| Data collection | RD-A |
| Magnification | x105,000 |
| Voltage (kV) | 300 |
| Electron Microscope | FEI Titan Krios G3i |
| Defocus (um) | -0.9 to -2.7 |
| Pixel size (Å) | 0.86 |
| Total dose (e^-^/ Å^2^) | 40.39 |
| Number of frames | 40 |
| Number of micrographs | 18 466 |
| Refinement |  |
| Number of total particles | 1 340 624 |
| GS-FSC Resolution (0.143, Å)^a^ | 2.55 |
| Model composition |  |
| Chains | 2 |
| Protein residues | 634 |
| r.m.s.d. |  |
| Bond lengths (Å) | 0.002 (0) |
| Bond angles (°) | 0.563 (0) |
| Validation |  |
| MolProbity score | 1.14 |
| Clash score | 3.49 |
| Ramachandran plot |  |
| Favored (%) | 98.57 |
| Allowed (%) | 1.43 |
| Disallowed (%) | 0.00 |
| CC Mask | 0.87 |
| ^a^Gold-Standard Fourier-Shell Correlation |  |

| **Extended Data Table 2. RD-21 cryo-EM data collection and refinement statistics.** | |
| --- | --- |
| Data collection | RD-21 |
| Magnification | x105,000 |
| Voltage (kV) | 300 |
| Electron Microscope | FEI Titan Krios G3i |
| Defocus (um) | -0.9 to -2.7 |
| Pixel size (Å) | 0.86 |
| Total dose (e^-^/ Å^2^) | 40.39 |
| Number of frames | 40 |
| Number of micrographs | 18 466 |
| Refinement |  |
| Number of total particles | 380,180 |
| GS-FSC Resolution (0.143, Å)^a^ | 2.64 |
| Model composition |  |
| Chains | 2 |
| Protein residues | 638 |
| r.m.s.d. |  |
| Bond lengths (Å) | 0.003 (0) |
| Bond angles (°) | 0.757 (1) |
| Validation |  |
| MolProbity score | 1.55 |
| Clash score | 4.16 |
| Ramachandran plot |  |
| Favored (%) | 97.63 |
| Allowed (%) | 2.37 |
| Disallowed (%) | 0.00 |
| CC Mask | 0.86 |
| ^a^Gold-Standard Fourier-Shell Correlation |  |

| **Extended Data Table 3. RD-23 cryo-EM data collection and refinement statistics.** | |
| --- | --- |
| Data collection | RD-23 |
| Magnification | x105,000 |
| Voltage (kV) | 300 |
| Electron Microscope | FEI Titan Krios G3i |
| Defocus (um) | -0.9 to -2.7 |
| Pixel size (Å) | 0.86 |
| Total dose (e^-^/ Å^2^) | 40.39 |
| Number of frames | 40 |
| Number of micrographs | 18 466 |
| Refinement |  |
| Number of total particles | 475,068 |
| GS-FSC Resolution (0.143, Å)^a^ | 2.66 |
| Model composition |  |
| Chains | 2 |
| Protein residues | 634 |
| r.m.s.d. |  |
| Bond lengths (Å) | 0.004 (0) |
| Bond angles (°) | 0.750 (0) |
| Validation |  |
| MolProbity score | 1.85 |
| Clash score | 2.49 |
| Ramachandran plot |  |
| Favored (%) | 96.19 |
| Allowed (%) | 3.49 |
| Disallowed (%) | 0.32 |
| CC Mask | 0.77 |
| ^a^Gold-Standard Fourier-Shell Correlation |  |

| **Extended Data Table 4. RD-25 cryo-EM data collection and refinement statistics.** | |
| --- | --- |
| Data collection | RD-25 |
| Magnification | x105,000 |
| Voltage (kV) | 300 |
| Electron Microscope | FEI Titan Krios G3i |
| Defocus (um) | -0.9 to -2.7 |
| Pixel size (Å) | 0.86 |
| Total dose (e^-^/ Å^2^) | 40.39 |
| Number of frames | 40 |
| Number of micrographs | 18 466 |
| Refinement |  |
| Number of total particles | 485,376 |
| GS-FSC Resolution (0.143, Å)^a^ | 2.58 |
| Model composition |  |
| Chains | 2 |
| Protein residues | 636 |
| r.m.s.d. |  |
| Bond lengths (Å) | 0.003 (0) |
| Bond angles (°) | 0.646 (0) |
| Validation |  |
| MolProbity score | 1.52 |
| Clash score | 2.78 |
| Ramachandran plot |  |
| Favored (%) | 97.31 |
| Allowed (%) | 2.53 |
| Disallowed (%) | 0.16 |
| CC Mask | 0.84 |
| ^a^Gold-Standard Fourier-Shell Correlation |  |

| **Extended Data Table 5. RF-21 cryo-EM data collection and refinement statistics.** | |
| --- | --- |
| Data collection | RF-21 |
| Magnification | x105,000 |
| Voltage (kV) | 300 |
| Electron Microscope | FEI Titan Krios G3i |
| Defocus (um) | -0.9 to -2.7 |
| Pixel size (Å) | 0.86 |
| Total dose (e^-^/ Å^2^) | 40.39 |
| Number of frames | 40 |
| Number of micrographs | 18 466 |
| Refinement |  |
| Number of total particles | 21,895 |
| GS-FSC Resolution (0.143, Å)^a^ | 3.17 |
| Model composition |  |
| Chains | 40 |
| Protein residues | 12,760 |
| r.m.s.d. |  |
| Bond lengths (Å) | 0.006 (0) |
| Bond angles (°) | 0.738 (174) |
| Validation |  |
| MolProbity score | 1.78 |
| Clash score | 8.08 |
| Ramachandran plot |  |
| Favored (%) | 95.09 |
| Allowed (%) | 4.91 |
| Disallowed (%) | 0.00 |
| CC Mask | 0.82 |
| ^a^Gold-Standard Fourier-Shell Correlation |  |

| **Extended Data Table 6. RF-23 cryo-EM data collection and refinement statistics.** | |
| --- | --- |
| Data collection | RF-23 |
| Magnification | x105,000 |
| Voltage (kV) | 300 |
| Electron Microscope | FEI Titan Krios G3i |
| Defocus (um) | -0.9 to -2.7 |
| Pixel size (Å) | 0.86 |
| Total dose (e^-^/ Å^2^) | 40.39 |
| Number of frames | 40 |
| Number of micrographs | 18 466 |
| Refinement |  |
| Number of total particles | 25,485 |
| GS-FSC Resolution (0.143, Å)^a^ | 3.20 |
| Model composition |  |
| Chains | 44 |
| Protein residues | 13,948 |
| r.m.s.d. |  |
| Bond lengths (Å) | 0.005 (0) |
| Bond angles (°) | 0.664 (247) |
| Validation |  |
| MolProbity score | 1.82 |
| Clash score | 7.52 |
| Ramachandran plot |  |
| Favored (%) | 93.98 |
| Allowed (%) | 5.75 |
| Disallowed (%) | 0.27 |
| CC Mask | 0.80 |
| ^a^Gold-Standard Fourier-Shell Correlation |  |

| **Extended Data Table 7. RF-25 cryo-EM data collection and refinement statistics.** | |
| --- | --- |
| Data collection | RF-25 |
| Magnification | x105,000 |
| Voltage (kV) | 300 |
| Electron Microscope | FEI Titan Krios G3i |
| Defocus (um) | -0.9 to -2.7 |
| Pixel size (Å) | 0.86 |
| Total dose (e^-^/ Å^2^) | 40.39 |
| Number of frames | 40 |
| Number of micrographs | 18 466 |
| Refinement |  |
| Number of total particles | 25,029 |
| GS-FSC Resolution (0.143, Å)^a^ | 3.18 |
| Model composition |  |
| Chains | 48 |
| Protein residues | 15,264 |
| r.m.s.d. |  |
| Bond lengths (Å) | 0.005 (0) |
| Bond angles (°) | 0.715 (18) |
| Validation |  |
| MolProbity score | 1.83 |
| Clash score | 8.19 |
| Ramachandran plot |  |
| Favored (%) | 94.26 |
| Allowed (%) | 5.49 |
| Disallowed (%) | 0.26 |
| CC Mask | 0.79 |
| ^a^Gold-Standard Fourier-Shell Correlation |  |

| **Extended Data Table 8. HF-23 cryo-EM data collection and refinement statistics.** | |
| --- | --- |
| Data collection | HF-23 |
| Magnification | x105,000 |
| Voltage (kV) | 300 |
| Electron Microscope | FEI Titan Krios G3i |
| Defocus (um) | -0.9 to -2.7 |
| Pixel size (Å) | 0.86 |
| Total dose (e^-^/ Å^2^) | 40.39 |
| Number of frames | 40 |
| Number of micrographs | 18 466 |
| Refinement |  |
| Number of total particles | 70,789 |
| GS-FSC Resolution (0.143, Å)^a^ | 2.91 |
| Model composition |  |
| Chains | 44 |
| Protein residues | 13,994 |
| r.m.s.d. |  |
| Bond lengths (Å) | 0.004 (0) |
| Bond angles (°) | 0.536 (173) |
| Validation |  |
| MolProbity score | 1.14 |
| Clash score | 2.56 |
| Ramachandran plot |  |
| Favored (%) | 97.50 |
| Allowed (%) | 2.50 |
| Disallowed (%) | 0.00 |
| CC Mask | 0.89 |
| ^a^Gold-Standard Fourier-Shell Correlation |  |

| **Extended Data Table 9. HF-25 cryo-EM data collection and refinement statistics.** | |
| --- | --- |
| Data collection | HF-25 |
| Magnification | x105,000 |
| Voltage (kV) | 300 |
| Electron Microscope | FEI Titan Krios G3i |
| Defocus (um) | -0.9 to -2.7 |
| Pixel size (Å) | 0.86 |
| Total dose (e^-^/ Å^2^) | 40.39 |
| Number of frames | 40 |
| Number of micrographs | 18 466 |
| Refinement |  |
| Number of total particles | 23,139 |
| GS-FSC Resolution (0.143, Å)^a^ | 3.08 |
| Model composition |  |
| Chains | 48 |
| Protein residues | 15,120 |
| r.m.s.d. |  |
| Bond lengths (Å) | 0.003 (0) |
| Bond angles (°) | 0.559 (38) |
| Validation |  |
| MolProbity score | 1.30 |
| Clash score | 3.91 |
| Ramachandran plot |  |
| Favored (%) | 97.40 |
| Allowed (%) | 2.60 |
| Disallowed (%) | 0.00 |
| CC Mask | 0.87 |
| ^a^Gold-Standard Fourier-Shell Correlation |  |
